# Werner syndrome helicase modulates G4 DNA-dependent transcription and opposes mechanistically distinct senescence-associated gene expression programs

**DOI:** 10.1101/023739

**Authors:** Weiliang Tang, Ana I. Robles, Richard P. Beyer, Lucas T. Gray, Giang Hong Nguyen, Junko Oshima, Nancy Maizels, Curtis C. Harris, Raymond J. Monnat

## Abstract

Werner syndrome (WS) is a prototypic heritable adult human progeroid syndrome in which signs of premature aging are associated with genetic instability and an elevated risk of specific types of cancer. We have quantified mRNA and microRNA (miRNA) expression in WS patient fibroblasts, and in WRN-depleted fibroblasts. Genes down-regulated in WS patient fibroblasts were highly enriched in G-quadruplex (G4) DNA motifs. The strength, location and strand specificity of this association provide strong experimental evidence that G4 motifs or motif-dependent G-quadruplexes are bound by WRN in human cells to modulate gene expression. The expression of many miRNAs was perturbed by loss of WRN function. WRN depletion altered the expression of >500 miRNAs. miRNAs linked to cell signaling, genome stability assurance and tumorigenesis were among the small number of these miRNAs that were persistently altered in WS patient fibroblasts. An unexpected and highly distinct finding in WS cells was the coordinate over-expression of nearly all cytoplasmic tRNA synthetases and their associated AIMP proteins. Our results provide new insight into WS pathogenesis, and identify therapeutically accessible mechanisms that may drive disease pathogenesis in WS and in the general population.

## Introduction

Werner syndrome (WS; OMIM #277700) is an autosomal recessive adult human progeroid syndrome caused by loss of function mutations in the *WRN* RECQ helicase gene (1) that encodes DNA-dependent ATPase, 3’- to -5’ helicase and 3’- to -5’ exonuclease activities (2, 3). The WS clinical phenotype resembles premature aging: graying and loss of hair, bilateral cataract formation and scleroderma-like skin changes appear in the second decade of life in conjunction with short stature. WS patients are at elevated risk for important, age-associated diseases such as atherosclerosis with myocardial infarction and stroke; cancer; osteoporosis; and diabetes mellitus. Of note, there is no elevated risk for Alzheimer disease or other dementias apart from those associated with vascular disease (4).

Cancer and cardiovascular disease are the leading causes of premature death in WS, most often in the fifth or sixth decades of life (4, 5). The cancer risk in WS patients is limited to a small number of specific neoplasms: acral lentiginous melanoma; meningiomas; soft tissue sarcomas; primary bone tumors, chiefly osteosarcomas; follicular thyroid carcinoma; and likely both leukemia and antecedent myelodysplasia (6). Loss of function mutations in two other members of the five member human RECQ helicase gene family, *BLM* and *RECQL4,* respectively cause the cancer predisposition syndromes Bloom syndrome (BS), and Rothmund-Thomson syndrome (RTS) and variants. Both BS and RTS have additional, associated developmental findings (2, 3).

Biochemical analyses have identified common biochemical substrates for both the WRN and BLM RECQ helicases. These substrates include forked or bubble DNAs, D-and T-loops, Holliday junctions and G-quadruplex structures (2, 3). G-quadruplexes are stable, non-B-form DNA structures that can be formed from nucleic acids containing G4 motifs of sequence G_≥3_N_1-x_G_≥3_N_1-x_G_≥3_N_1-x_G_≥3_. G4 motifs are highly enriched in rDNA, near transcription start sites (TSSs), at the 5’ ends of first introns and in exons and introns of many oncogenes (7). We showed previously that G4 motifs are regulatory targets for the BLM RECQ helicase (8), and for binding by the xeroderma pigmentosum-related helicases XPB and XPD (9). Here we quantify mRNA and miRNA expression in WS patient and WRN-depleted primary fibroblasts. Genes and miRNAs whose expression is modulated by WRN mutation or loss identify potential mechanisms by which WRN may modulate gene expression, and functional pathways that may drive WS pathogenesis and disease risk.

## Results

### Expression profiling identifies WRN-regulated genes and miRNAs

We quantified and compared mRNA and miRNA expression in WS patient (WS) versus matched control (NM) donor primary fibroblasts, and in isogenic control primary human fibroblasts depleted of WRN (**Table S1, Fig. S1, Fig. S2**). Our choice of fibroblasts was driven by the prominent cutaneous features of WS (4, 10), and by the well-defined WS fibroblast cellular phenotype (11). We used array-based profiling to allow a direct comparison with prior attempts to identify altered gene expression in WS patient or WRN altered cells, nearly all of which used comparable expression analysis platforms (12–15)

Principal component analysis (PCA) of expression data showed sample partitioning by genotype for mRNA and miRNA expression, with no additional structure reflecting patient/donor age, gender or passage number (Fig. 1). Significant differential mRNA expression, defined by an absolute fold difference of ≥1.5 with an FDR of <0.05 versus matched controls, was observed in WS fibroblasts for 628 mRNAs with increased expression and 612 mRNAs with decreased expression (**Table S2**). These cut-offs are identical to our previous analysis of mRNA in BS patient and BLM-depleted cells (8), and were chosen after a sensitivity analysis using different cut-off values (see **Supporting Information** Methods for additional detail). WRN-depleted fibroblasts displayed 839 mRNAs with significant increased expression and 674 mRNAs with significant decreased expression *versus* nonspecific (NS)-shRNA-depleted controls (**Table S3**). A total of 281 (or 11.4%) of these mRNAs were common to both WS cells and WRN-depleted fibroblasts, with 198 of these (70%) showing coordinate increased or decreased expression in both cell types (Fig. 1; **Table S4**).

**Figure 1.**
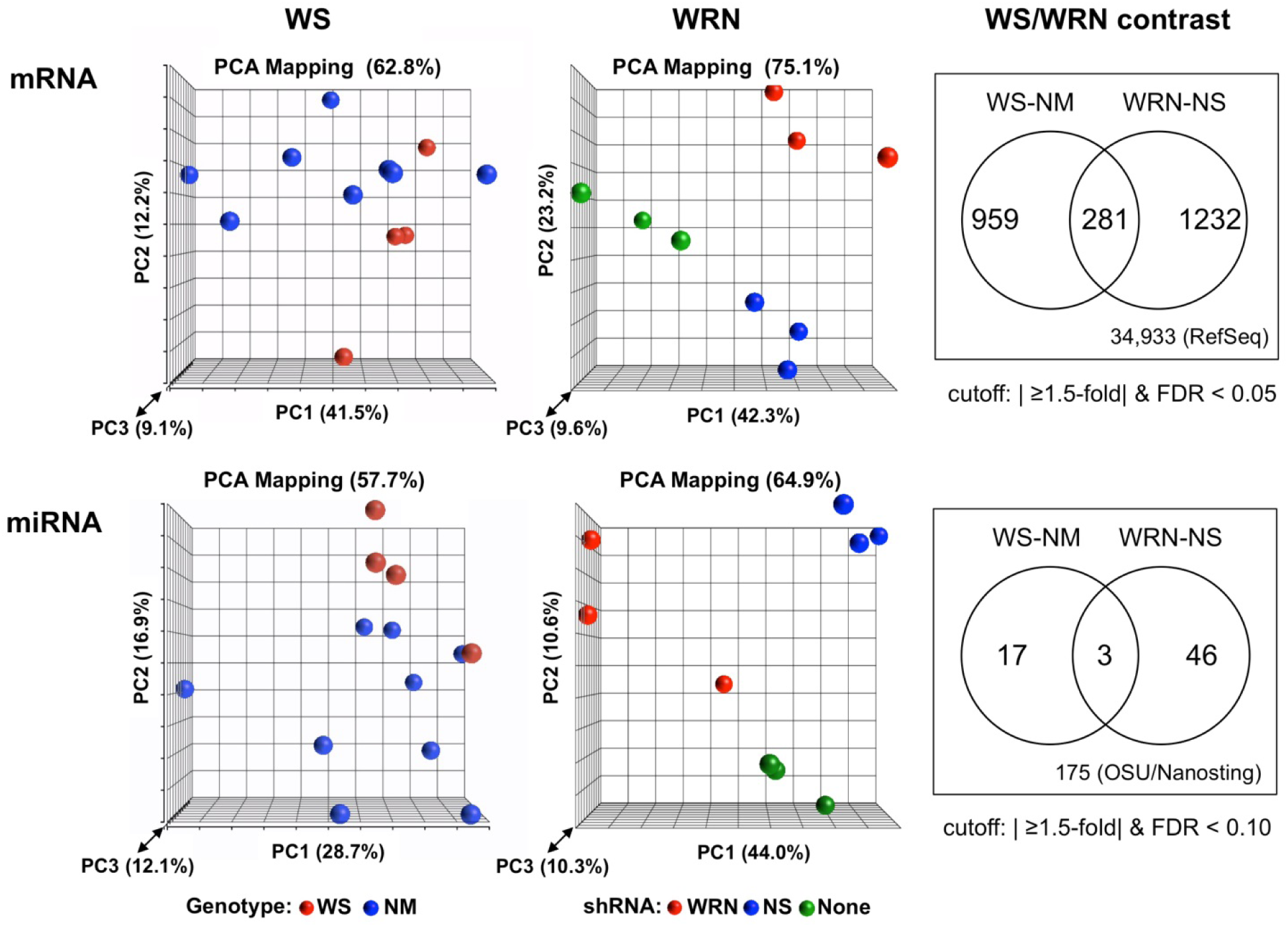
mRNA and miRNA expression patterns distinguish WS and WRN-depleted cells from comparable controls. Principal component analyses (PCA) of mRNA expression (top row) and miRNA expression (bottom row) in Werner syndrome patient (WS, red) and matched control donor (NM, blue) cells, and in WRN-depleted (WRN, red), nonspecific scrambled shRNA treated (NS, blue), and untreated (None, green) cells. Venn diagrams (last column) depict genes (top) and miRNAs (bottom) with significantly altered expression in WS patient or WRN-depleted human primary fibroblasts.

Differentially expressed miRNAs were identified using criteria identical to our previous analysis of miRNA expression in BS patient and BLM-depleted cells (8): by an absolute fold difference of ≥1.5 and a FDR of <0.10 versus controls. Nineteen miRNAs were differentially expressed in WS *versus* control fibroblasts, and 92 miRNAs in WRN-depleted versus NS-depleted fibroblasts. Ten miRNAs were differentially expressed in WS and in WRN-depleted cells: four displayed coordinate increased expression in both cell types (miR-323a-5p, and miR-632, 636 and 639), while the remaining six displayed increased expression in WS cells and decreased expression in WRN-depleted cells (Fig. 1; **Table S5a**).

We also examined expression within each cell type of all miRNAs that could be analyzed using each expression platform. WS cells displayed 23 different miRNAs that were differentially expressed versus matched control donors by the above criteria, with 15 having increased expression and 8 reduced expression (**Table S5b**). A corresponding analysis of WRN-depleted versus NS-depleted cells revealed 543 miRNAs significantly altered using the above criteria, with the vast majority (522 of 543 miRNAs or 96%) showing reduced expression)(**Table S5c**). Perturbed miRNA expression in WRN-depleted cells was remarkably robust, with nearly 2/3 of these miRNAs (328 of 334 or 98%) displaying significant differential expression when we used an absolute fold difference of ≥2.0 and a FDR of <0.05 as cutoffs (**Table S5d**).

We used two approaches to determine whether altered miRNA expression might be driving altered mRNA expression and disease-associated pathways in WS patient fibroblasts. First, an experimentally-validated miRNA target genes list was used to determine whether a disproportionate number of miRNA-specific target genes were coordinately and reciprocally altered in expression with their cognate, targeting miRNA (**Table S6**). This analysis identified several miRNAs that might drive differential mRNA expression in WS and/or WRN-depleted cells (**Table S6a**). We also performed a more broadly based search for miRNA-target gene pairs as part of an IPA analysis of miRNA expression. This analysis used miRNA-target pairs from miTarBase, TargetScan and additional experimentally-observed miRNA-target pairs to identify miRNAs and target genes whose expression varied reciprocally, and identified 18 miRNAs and 218 target genes linked to specific pathways that may contribute to WS disease pathogenesis or disease risk (**Table S7**).

### G4 DNA motifs correlated with WRN-dependent differential gene expression

In order to determine whether G4 DNA motifs were correlated with WRN-dependent differential gene expression, we determined the frequency of G4 motifs near the transcription start site (TSS) and 5’ end of the first intron of genes differentially expressed in WS and/or WRN-depleted cells. An empirical FDR was calculated by comparing G4 motif frequencies in a set of comparably-sized control gene sets randomly selected from all expressed genes (Fig. 2).

**Fig. 2.**
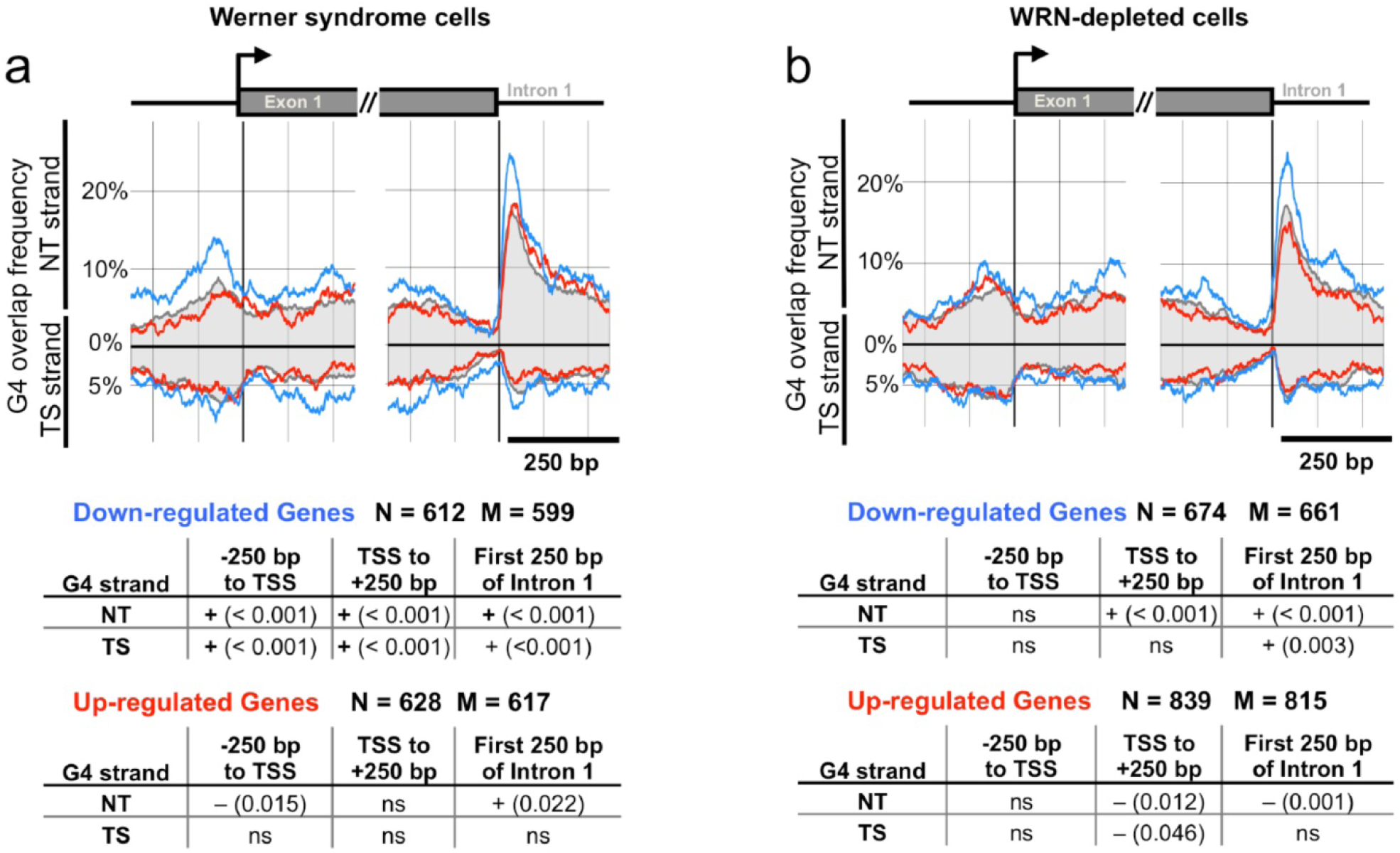
Enrichment of G4 DNA motifs near TSSs and exon 1/intron 1 boundaries of genes exhibiting differential expression in WS or WRN-depleted human fibroblasts. G4 motif frequencies as a function of location in genes exhibiting differential regulation in (a) WS patient cells or in (b) WRN-depleted control fibroblasts. G4 motif counts relative to the transcription start site (TSS) are shown for the non-transcribed (NT) and transcribed (TS) strand of genes with decreased (blue line) or increased (red line) expression, versus average G4 motif frequencies (gray shaded area) for all genes surveyed. Bottom panel indicates FDRs of observed G4 motif frequencies versus randomly selected control gene sets. N = gene accessions in DEG table; M = number of matching TSS locations from gene location tables; ns: non-significant (FDR>0.05).

Genes exhibiting reduced expression in WS fibroblasts were highly enriched in G4 motifs on both DNA strands up-and downstream of the TSS, and at the 5’ end of first intron (FDR<0.001). G4 motifs were correlated with increased gene expression in WS cells only at the 5’ end of intron 1 (FDR=0.022), and with depletion of G4 motifs in the nontranscribed (NT) strand in the 250 bp upstream of the TSS (FDR=0.015)(**Tables S8, S9**).

Genes exhibiting reduced expression in WRN-depleted cells were significantly enriched for G4 motifs on the NT strand downstream of the TSS (FDR<0.001), and on both strands at the 5’ end of intron 1 (TS, FDR=0.003; NT, FDR<0.001)(Fig. 2). Conversely, genes whose expression was increased by WRN depletion lacked G4 motifs on either strand downstream of the TSS (FDR = 0.012), or on the NT strand at the 5’ end of intron 1 (FDR < 0.001)(**Tables S10, S11**). These results indicate that G4 DNA motifs near the TSS and at the 5’ end of first introns may be physiologic targets for WRN binding leading to altered gene expression in human cells.

### Molecular functions and pathways altered in WS and WRN-depleted cells

Ingenuity Pathway Analysis (IPA Ingenuity Systems; www.ingenuity.com) and gene set enrichment analysis (GSEA) were used to identify molecular and cellular functions, canonical pathways and networks perturbed in WS patient and WRN-depleted primary fibroblasts (Table 1, **Tables S12-S14**). Top-ranked functions perturbed in both WS and WRN-depleted cells were ‘cell growth/proliferation’, ‘cell death/survival’ and ‘cell cycle’. ‘Cellular development’ and ‘cell movement’ were highly ranked in WS cells, as were ‘cellular assembly/organization’ and ‘DNA replication/recombination/repair’ in WRN-depleted cells. Four top-ranked canonical pathways altered in WS fibroblasts involved protein synthesis: tRNA charging, and EIF2, mTOR and ‘regulation of eIF4 and p70/S6 kinase’ signaling. Signaling pathways represented 8 and 7 of the top 10 perturbed pathways in, respectively, WS patient and WRN-depleted fibroblasts with ‘aryl hydrocarbon receptor (AHR) signaling’ top-ranked in WRN-depleted cells and fifth-ranked in WS patient fibroblasts (Table 1, **Tables S12, S13**). The top-ranked perturbed networks in WS cells were focused on connective tissue biology and disease. These were of considerable interest given the strong cutaneous phenotype and increased risk of connective tissue neoplasms of WS (**Table S12**)(6, 10).

**Table 1.** Ingenuity Pathways Analysis (IPA) summary

| **Werner syndrome (WS) fibroblasts** |  |  | **WRN-depleted control fibroblasts** |  |  |
| --- | --- | --- | --- | --- | --- |
| **Top molecular/cellular functions** |  |  |  |  |  |
| ***Function*** | ***p value*** | ***molecules*** | ***Function*** | ***p value*** | ***molecules*** |
| cellular growth/proliferation | 3.8E-12 | 468 | cell cycle | 1.4E-21 | 260 |
| cell death/survival | 7.2E-11 | 408 | cellular growth/proliferation | 3.3E-21 | 494 |
| cellular development | 2.0E-09 | 377 | cell death/survival | 1.8E-19 | 472 |
| cell movement | 3.5E-07 | 287 | cellular assembly/organization | 1.1E-18 | 277 |
| cell cycle | 4.1E-07 | 180 | DNA replication/recomb/repair | 1.1E-18 | 176 |
| **Top canonical pathways** |  |  |  |  |  |
| ***Pathway*** | ***p value*** | ***ratio*** | ***Pathway*** | ***p value*** | ***ratio*** |
| tRNA charging | 1.26E-07 | 14/35 | AHR signaling | 3.10E-08 | 30/140 |
| EIF2 signaling | 1.35E-04 | 22/121 | cell cycle:G2/M reg | 4.38E-06 | 14/49 |
| integrin signaling | 1.66E-04 | 29/183 | polo-like kinase mitotic roles | 9.65E-05 | 16/66 |
| JAK/TYK in IFN signaling | 2.14E-04 | 8/23 | gap junction signaling | 2.90E-05 | 26/155 |
| AHR signaling | 3.09E-04 | 22/128 | IL6 signaling | 5.29E-05 | 21/116 |

Gene set enrichment analysis (GSEA) identified gene sets that contained disproportionate numbers of genes that were differentially expressed in WS and/or WRN-depleted cells (Table 2, **Table S14**). Top-ranked C2_CP (Canonical Pathways) gene sets identified by this analysis for WS cells were Reactome tRNA/cytosolic tRNA aminoacylation and KEGG Aminoacyl tRNA biosynthesis (**Table S15**). C3_TFT (Transcriptional Factor Target) gene sets were significantly enriched for 62 binding sites encompassing 66 different TFs in differentially expressed genes WS cells, and for 26 binding sites encompassing 26 TFs in WRN-depleted cells. Binding sites for 7 TFs were enriched in both WS and WRN-depleted cells, together with differential expression of 5 TFs in WS cells and 2 TFs in WRN-depleted cells (**Table S16**). Our G4 motif, IPA and GSEA analysis results collectively suggest that WRN modulates gene expression via a regulatory network that includes G4 DNA elements and TFTs in *cis*, and miRNAs and TFs in *trans*.

**Table 2.**
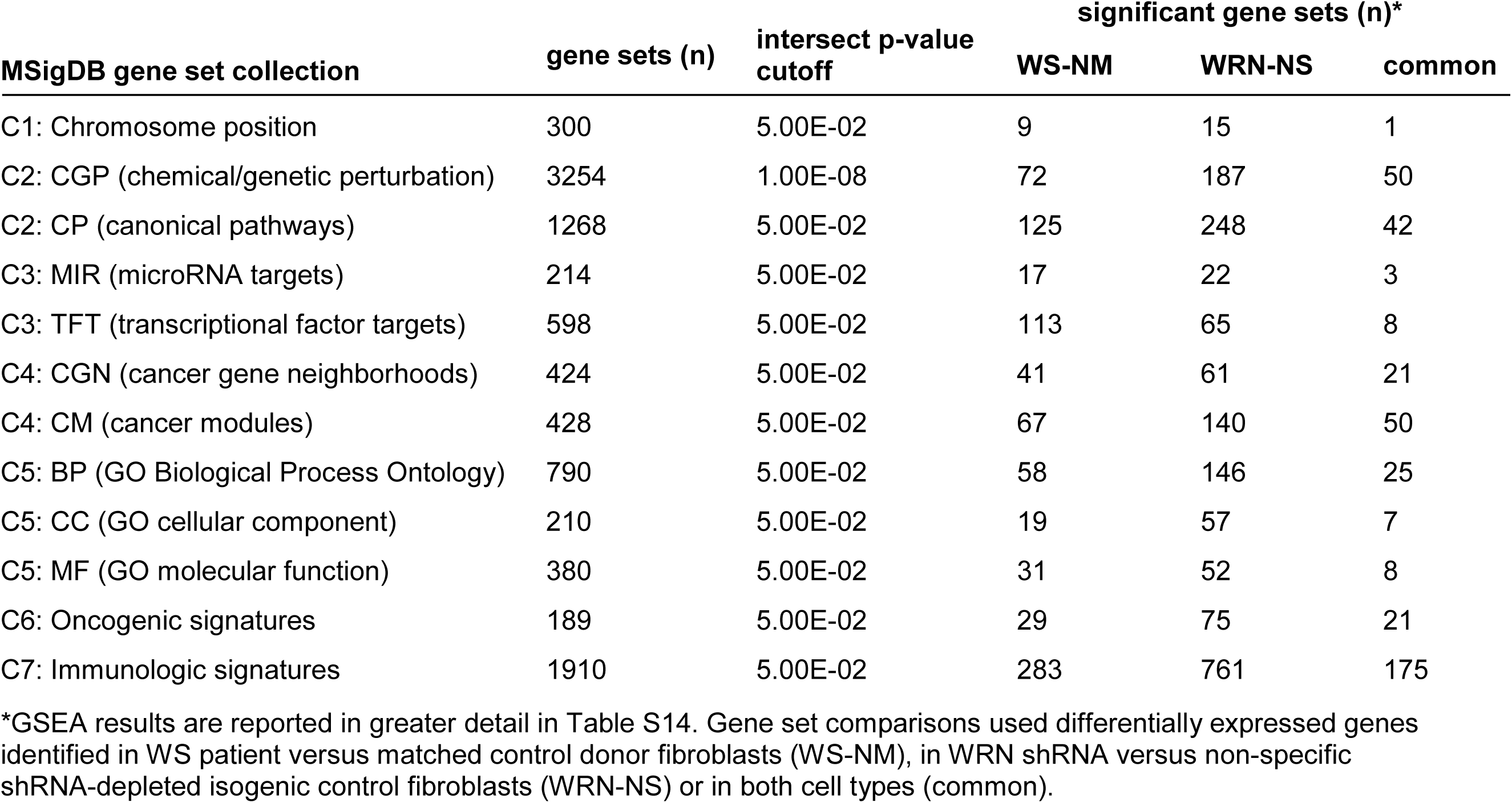
Gene Set Enrichment Analysis (GSEA) summary.

### Coordinate upregulation of cytoplasmic tRNA synthetase and AIMP genes in WS

A surprising finding in IPA and GSEA analyses of WS cells was ‘tRNA charging’ as a top-ranked altered canonical pathway (IPA p-value=1.26E-07) in conjunction with several other pathways that regulate protein synthesis (Table 1, **Table S12**). The tRNA charging pathway consists of genes encoding 19 cytoplasmic and 20 mitochondrial tRNA synthetases (ARSs), 3 ARS-interacting multifunctional proteins (AIMPs) and several phosphatases (Reactome tRNA aminoacylation pathway REACT_15482.3). WS cells displayed significantly increased expression of 15 of the 18 cytoplasmic ARS genes and the 2 AIMP genes included on our array platform (Fig. 3). Of note, ‘tRNA charging’ did not appear among the 100 top-ranked pathways in WRN-depleted cells (IPA p-value=0.29), which displayed increased expression of only 3 cytoplasmic ARS genes and AIMP1 in conjunction with reduced expression of 2 ARS genes (Fig. 3, **Table S12**). This pattern of coordinate increased cytoplasmic ARS/AIMP expression was both highly distinctive and statistically significant: WS cells were the top-ranked data set for this expression pattern among 5,474 comparison samples from GSE GEO (Gene Expression Omnibus) data (Fig. 3, **Table S17**). Of note, ARS/AIMP expression was not significantly perturbed in recently described human *WRN* knockout embryonal stem cells (ESCs) or in differentiated mesenchymal stem cells (MSCs)(16).

**Fig. 3.**
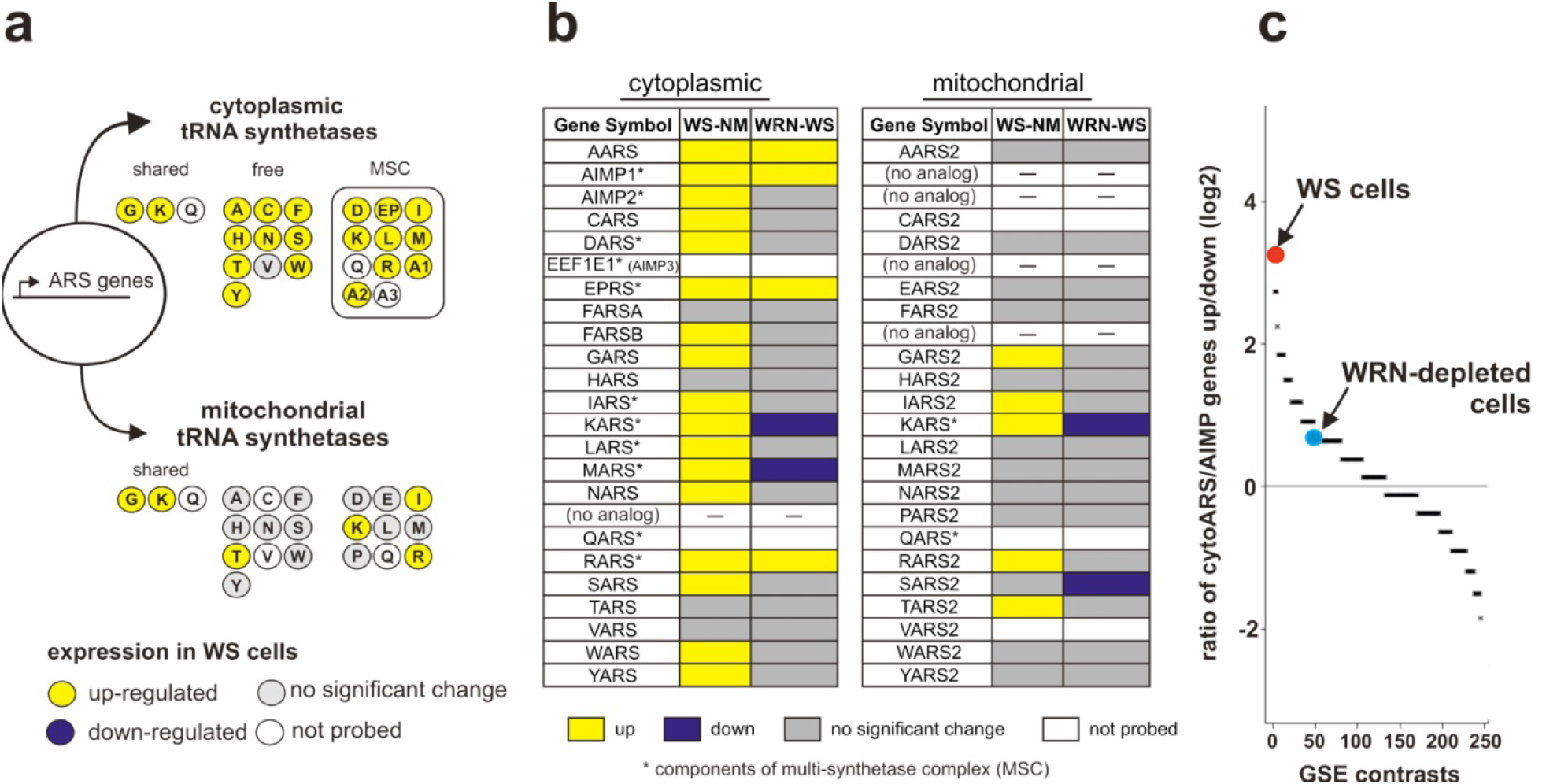
WS cells display increased expression of many cytoplasmic tRNA synthetase and AIMP genes. (a) tRNA synthetase and AIMP proteins are nuclear-encoded and partitioned to the cytoplasm or mitochondrion with three shared (GARS, KARS and QARS). Eleven ARS+AIMP proteins form a cytoplasmic multisynthase complex (MSC). Individual proteins are represented by circles with single letter amino acid or AIMP number labels. Shading indicates expression in WS cells. (b) ARS/AIMP expression in WS (WS-NM) and WRN-depleted (WRN-NS) cells. (c) ARS+AIMP upregulation ratio in WS and WRN-depleted cells versus 56 GEO Series (GSE) data sets (see *SI Materials and Methods* for additional detail).

### WRN suppresses mechanistically distinct senescence programs

Cellular senescence is a prominent phenotype in both WS patient and WRN-depleted cells (11). We thus determined whether senescence-associated genes and gene sets representing mechanistically distinct senescence pathways were differentially expressed in WS and/or WRN-depleted cells (**Tables S18, S19**). Six gene sets were used: a senescence gene ‘superset’ (156 genes; REACT_169274.2); and gene sets reflecting oncogene-induced senescence (30 genes; REACT_169325.2); oxidative stress-induced senescence (89 genes; REACT_169436.2); DNA damage/telomere stress-induced senescence (58 genes; REACT_169185.2); the senescence-associated secretory phenotype (SASP)(75 genes; REACT_169168.2); and ‘SASP_components_genes’, 81 genes encoding cellular or secreted SASP factors assembled from (17, 18)).

GSEA analysis indicated significant enrichment of genes that were differentially expressed in WS and/or WRN-depleted cells in four of these gene sets. Increased gene expression was observed for differentially expressed genes in the ‘DNA damage/telomere-induced senescence’ gene set (in WS cells), and the ‘SASP_components_genes’ set (in WRN-depleted cells), with most other genes displaying reduced expression (Fig. 4). These results indicate that *WRN* mutation and WRN depletion significantly alter the expression of many genes associated with mechanistically distinct, senescence-associated gene expression programs. Several different senescence-associated pathways included in this analysis, most notably the senescence-associated secretory phenotype (SASP) pathway, may serve as potential drivers of WS cellular and organismal phenotypes.

**Fig. 4.**
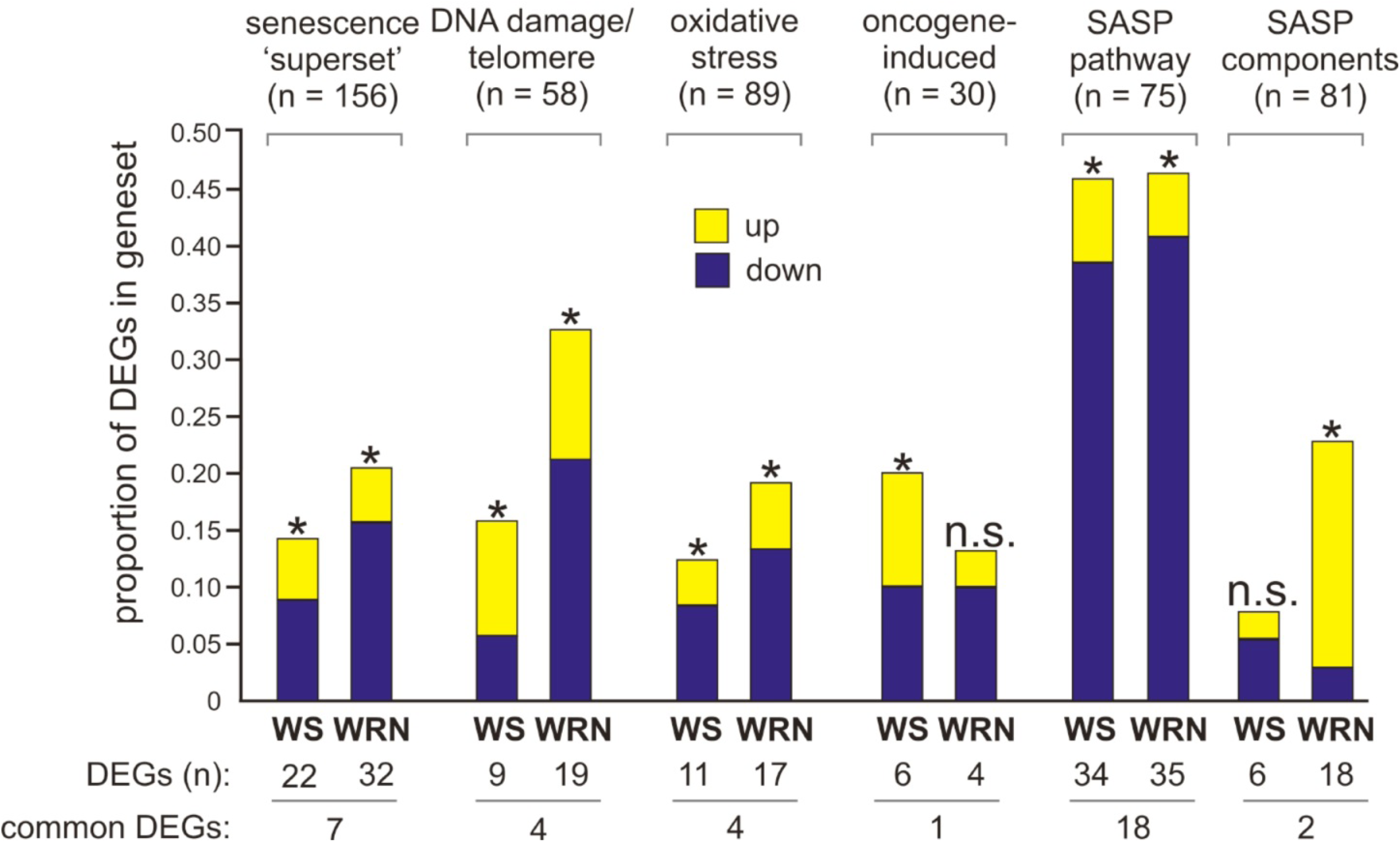
WRN loss perturbs the expression of mechanistically distinct, senescence-associated genes and gene sets. Bar graphs indicate proportion of differentially expressed genes (DEGs) in senescence-associated gene sets in WS patient (WS) or WRN-depleted (WRN) cells. Gene sets and gene number/set are indicated over bars, with shading indicating increased (yellow) or decreased (blue) expression in WS or WRN-depleted fibroblasts. GSEA intersect p values of <0.05 are indicated by asterisks. Common DEGs were regulated in the same direction across all 6 gene sets.

We also looked for significant enrichment for differentially expressed genes (DEGs) in gene expression data recently reported for human *WRN* mutant ESCs and MSCs (16). As anticipated, *WRN* mutant ESCs showed a very low proportion of DEGs across all six senescence-associated gene sets. *WRN* mutant MSCs most closely paralleled WRN-depleted, as opposed to WS patient, cells: MSCs displayed a similar distribution, though lower proportion, of DEGs together with a higher ratio of genes with reduced as opposed to increased expression when compared to WS and WRN-depleted cells (**Fig. S6**).

### WRN and BLM regulate distinct sets of target genes

The WRN and BLM human RECQ helicases share many functional similarities including G4 motif-dependent gene expression. We thus compared gene expression changes in WS cells with cells from patients with Bloom syndrome (BS). Among 2,179 genes differentially expressed in WS and BS cells, >90% were perturbed only in WS or in BS cells (**Fig. S3**)(8). Among the 214 genes differentially expressed in both WS and BS cells, a high fraction (87.4%) displayed coordinate increased or decreased expression and represented a wide range of different functions and pathways (**Table S20**). Common genes exhibiting reduced expression were enriched for G4 motifs on both strands upstream of the TSS (FDR=0.009 and 0.002), and on the NT strand at the 5’ end of intron 1 (FDR=0.041). In contrast, genes exhibiting increased expression were enriched in G4 motifs only on the NT strand, upstream of the TSS (FDR=0.025)(**Fig. S4**). A similar pattern was observed in WRN-or BLM-depleted cells: 75% of 2,282 differentially expressed genes were altered in only one cell type, and a high proportion of 563 common genes (93.1%) displayed coordinate increased or decreased expression (**Table S21, Fig. S5**). Genes displaying coordinate decreased expression were enriched in functions that can be tied directly to chromosome structure, DNA metabolism and genome stability maintenance: histone, centromeric protein, cell division cycle (CDC/CDK), mitotic division, replication and repair genes including MCMs, POLD, RAD51 and multiple Fanconi anemia genes)(**Table S21**). A majority of differentially expressed miRNAs in WS or BS cells were unique to one cell type: six of 52 miRNAs shared between WS and/or BS cells were coordinately expressed, as were 10 of 58 miRNAs in WRN-and BLM-depleted cells (**Table S22**). These results collectively indicate that WS and BS cells have largely distinct patterns of gene and miRNA expression, though share small subsets of genes that are enriched for G4 motifs and coordinately regulated miRNAs.

## Discussion

We used RNA expression profiling to identify genes and miRNAs that were differentially expressed in cells that lacked WRN RECQ helicase protein function as a result of *WRN* mutations or WRN protein depletion. This experimental design was chosen to unambiguously identify genes and miRNAs whose expression was altered in a WRN-dependent manner; to identify potential mechanisms by which WRN modulates gene expression; to reveal adaptations in gene and miRNA expression in response to acute or prolonged loss of WRN function; and to identify functional pathways that might drive WS pathogenesis and disease risk. This design thus substantially extended previous analyses of gene expression in WS cells (12-15), and in recently reported WRN-mutant ESC and MSC cultures where we have further analyzed reported results (16).

### G-quadruplexes are targets for regulation by WRN

Biochemical evidence has long suggested G-quadruplex structures may be targets for the RECQ helicases WRN and BLM. Consistent with this idea, we previously showed that the Bloom syndrome RECQ helicase (and the TFIIH XPB and XPD disease-associated helicases) also modulate transcription in a G4 motif-dependent manner (8, 9). In order to test the hypothesis that WRN may modulate gene expression in human cells in a G4 motif-dependent manner, we determined whether genes with reduced expression in WS cells were highly enriched in G4 motifs both upstream of the transcription start site (TSS), near the exon 1/intron 1 boundary and in the first 250 bp of intron 1 on one or both DNA strands (Fig. 2). These G4 motifs are well-positioned to influence gene expression by altering transcription initiation, RNA Pol II pausing, and pre-mRNA splicing.

Our results collectively provide strong evidence that G4 motifs and/or G-quadruplex structures formed from G4 motifs are physiologic targets for transcription regulation in human cells. A simple regulatory model for WRN G4-dependent modulation of gene expression involves WRN binding and unwinding of G-quadruplex structures *in vivo* to facilitate the transcription of G4 motif-rich genes. Further analyses of the G4 motifs in genes differentially expressed in WS, BS or XPB/XPD cells should provide additional insight into G4 motif and G-quadruplex structures that are targeted by specific helicases in human cells.

### miRNA expression is strongly perturbed in WRN-depleted cells

WRN is also able to modulate the expression of a remarkably large number of miRNAs. For example, in WRN-depleted cells 543 miRNAs identified using previously published criteria (8) were significantly differentially expressed in WRN-depleted versus NS control cells, with the vast majority of these (96%) displaying reduced expression. Among the small number of persistently altered miRNAs in WS cells, several were identified that may play roles in WS disease pathogenesis (**Tables S5a-d**). For example, over-expression of the miR-181 family have been implicated in leukemogenesis (19), whereas down-regulation of miR-632 and 636 have been implicated in myelodysplasia (20). Further analyses of miRNA-target gene pairs also identified genes and canonical pathways altered in conjunction with persistent miRNA expression that may drive WS disease pathogenesis as we had previously observed for Bloom syndrome (8).

### WRN regulates disease-associated functional pathways

*WRN* mutation or WRN depletion both altered the expression of genes involved in cell cycle control, growth and proliferation; DNA replication, recombination and repair; and DNA damage checkpoint signaling (Table 1, **Tables S2-4**). These pathways have been mechanistically linked to genomic instability, cell proliferation defects and cellular or organismal senescence (17, 21).

WRN also modulates several overlapping, mechanistically distinct senescence-associated gene expression programs. This differs by pathway and cell type (WS versus WRN-depleted; Fig. 4), and our analysis identified gene expression changes that could drive both intrinsic, chromatin state-associated as well as secretory aspects of aging. Of note, these senescence-associated gene expression programs were not as strongly perturbed in recently reported WRN mutant human ESC and MSC cells that have been proposed as WS disease models (**Figure S6**)(16). It will be important to characterize the stability and phenotype of WRN-mutant MSCs, and to examine gene and miRNA expression changes in this and other cellular models of WS to determine whether they provide good experimental systems to use to investigate cellular, cell lineage-and tissue-specific consequences of WRN loss of function.

### Coordinate over-expression of tRNA charging genes in WS cells

A surprising and unexpected finding in our mRNA expression analyses of WS cells was the coordinate increased expression of nearly all of the cytoplasmic tRNA synthetases (ARSs) and their associated interacting proteins (AIMPs). This gene expression signature was highly distinctive and statistically highly significant, and provides new insight into WS disease pathogenesis mechanisms.

WS patient fibroblasts were the top-ranked data set for coordinate increased ARS+AIMP expression among 5,474 samples in 56 GEO Series (GSE) data sets that included a wide range of cancer, non-neoplastic disease and control samples (**Table S17; Fig. 3 and Table S12**). The mechanism(s) responsible for coordinate increased expression remain to be identified: ARS/AIMP expression does not appear to be strongly driven by G4 DNA motif depletion, or by MYC overexpression, which itself requires WRN expression to avoid rapid cellular senescence (22). MYC may, however, be indirectly influencing ARS+AIMP upregulation as we identified activation of the MYC transcription network in WS cells (see **Table S14**). We did identify differential enrichment for specific transcription factor target sites (TFTs) in WS versus WRN-depleted cells (**Table S16**). Further analysis of these TFTs may provide a more detailed understanding of the regulatory network that governs coordinate ARS and AIMP expression in WS cells, and that coordinately regulates ARS/AIMP expression in normal human cells.

The coordinate upregulation of ARS+AIMP genes in WS patient fibroblasts provides intriguing new insight into WS disease pathogenesis mechanisms. The cytoplasmic ARS and AIMP proteins are abundant, comparatively long-lived proteins with copy numbers ranging from 10^5^ to >10^6^/cell (23, 24). The increased expression of ARS and AIMP genes in WS cells likely represents a parallel increase in corresponding protein copy numbers in WS cells (25–27). Higher steady-state ARS and AIMP copy numbers could, together with altered EIF2, mTOR, and eIF4/S6 kinase pathway signaling, perturb global protein dynamics in WS cells (Table 1). Recent work consistent with the idea of altered global protein dynamics in WS include the observation that rapamycin and hydrogen sulfide can reduce protein aggregation, suppress oxidative stress and reduce DNA damage in WS cells (28) and in WRN-depleted fibroblasts (29). Altered translational fidelity, originally analyzed in WS fibroblasts with inconclusive results (see, e.g., (30)), could also be perturbed in WS cells by increased ARS+AIMP expression, persistent stress signaling and altered senescence-associated gene expression (31, 32).

The coordinate, increased expression of cytoplasmic tRNA synthetases in WS could also disrupt ‘mitonuclear’ balance between cytoplasmic and mitochondrial protein synthesis. Few genes are transcribed and translated in the mitochondrion. Despite this, balanced mitonuclear protein synthesis is clearly important for maintaining mitochondrial function and appears to be actively maintained by a regulatory network involving oxidative phosphorylation status, the mitochondrial unfolded protein response (UPR^mt^) and NAD+/Sir2/Foxo-dependent signaling. This system can also, in some systems, modulate longevity and senescence (33, 34).

A third, intriguing mechanism by which increased ARS and AIMP expression could promote WS disease pathogenesis is via ‘non-canonical’ functions in addition to tRNA charging. These non-canonical functions involve specific ARS and AIMP proteins, ARS protein splice isoforms (35) or proteolytic cleavage fragments, and modulate multiple disease-related metabolic, developmental, angiogenic, tumorigenic, immune and inflammatory pathways (36). The coordinate upregulation of many ARS and AIMP genes and their associated non-canonical functions provides a mechanistically appealing—if provocative—explanation for the pleiotropic organismal phenotype of WS.

### Potential therapeutic implications

Further testing the mechanistic ideas outlined above should advance our understanding of WS disease pathogenesis, and may reveal therapeutically accessible pathways that could be targeted both in WS patients and the general population. Global protein dynamics and translational fidelity in WS could be profitably reinvestigated using sensitive, quantitative global profiling assays (25–27, 31, 37, 38). Altered protein dynamics and/or mitonuclear balance, if identified, could be targeted with rapamycin (39) or resveratrol (40). Additional small molecules could also be used to increase NAD+ levels (e.g., PARP-1 inhibitors, nicotinamide and nicotinamide riboside)(41); induce the UPR^mt^ (doxycycline (33)); or activate SIRT1 (e.g., resveratrol and other small molecule activators (40)). Resveratrol itself may act via both SIRT1 and tRNA synthetase/PARP-1 pathways (40, 42). Many of these agents have immediate therapeutic potential, as they are or can be used in the clinic, and thus could be useful both in the care of WS patients and in the modulation of age-associated disease pathways revealed by and shared in common between WS and normal aging.

### Summary

Werner syndrome, the canonical human adult progeroid syndrome, is a remarkable example of a Mendelian disease that perturbs many aging and age-associated disease pathways (21). Our results provide the first systematic analysis of miRNA expression in WS, and substantially extend prior analyses of gene expression in WS cells (12–15) and in human stem cell models (16). We identified the outline of a WRN-dependent gene expression regulatory network that includes G4 DNA motifs and transcription factor target sites (TFTs) in *cis,* together with miRNAs and TFs in *trans.* Functional pathways linked by this network provide new insight into WS pathogenesis, and may identify new ways to modulate aging and age-associated disease risk in WS patients and potentially in normal individuals (18, 21).

## Materials and Methods (see also Methods in *Supporting Information* for additional detail and methods references)

### Cells

Primary human skin fibroblast strains (**Table S1**) from 4 mutation-typed WS patients (mean age 33.5 years, median age 33 years) or from 10 age-and gender-matched control donors (mean age 30.2 years, median age 28 years) were obtained from the Coriell Cell Repositories (Camden, NJ, USA). Control human primary fibroblast strain 82-6 was isolated as previously described (8), and WRN was depleted by lentiviral *WRN*-specific shRNA expression and verified by Western blot (**Figs. S1, S2**).

### mRNA and miRNA expression profiling

Whole genome transcript exon profiling and miRNA profiling were performed on Affymetrix GeneChip Human Exon 1.0 ST (Affymetrix, Santa Clara, CA) arrays and a custom miRNA microarray or Nanostring nCounter, and differentially expressed genes and miRNAs identified as previously described (8). Expression profiling data are archived under GEO accession number GSE62877.

### RNA expression analyses

Principal component analysis (PCA), Ingenuity pathway and gene set enrichment analyses were all performed as previously described (8). Gene Set Enrichment Analyses (GSEA) used all C1-C7 gene sets in the Molecular Signature Database version 4.0. miRTarBase was used to identify experimentally validated gene targets of miRNAs differentially expressed in WS and WRN-depleted cells as previously described (8), and differentially expressed miRNAs were used to seed IPA analysis of miRNA targets and pathways. G4 motifs were located in hg19 using Quadparser, then analyzed for their association with genes differentially regulated in WS or WRN-depleted cells as previously described (8). GSEA was used to determine whether differentially expressed genes were enriched in 6 senescence-associated gene sets (**Figure S6**, **Tables S18 and S19**), and to analyze the uniqueness and statistical significance of coordinate, increased ARS and AIMP expression in 5,474 samples contained in 56 GEO Series (GSE) data sets. GSEA p-values were calculated for three alternative expression hypotheses (up, down, or mixed).

## Acknowledgments

We thank the staffs at the Laboratory of Molecular Technology (NCI-Frederick), the Center for Ecogenetics and Environmental Health Functional Genomics and Proteomics Core (UW) and the Microarray Core Facility at the Ohio State University for microarray hybridization and scanning. Kensuke Kumamoto MD PhD and Xin Wei Wang PhD provided useful early comments on study design and implementation. Alden Hackmann provided graphics support, and Matt Kaeberlein, David Morris, Peter Rabinovitch and Susan Martinis for useful discussion. This work was supported by National Cancer Institute of the NIH award P01CA77852 to RJM, Jr.; by the Intramural Research Program of the National Cancer Institute, Center for Cancer Research, National Institutes of Health (NIH); by the National Institute of Environmental Health Sciences of the National Institutes of Health under Award Number 5P30ES007033; by R24AG042328 (International Registry of Werner Syndome) to JO; and by the NIH-Oxford MD/DPhil Fellowship Program (GHN). Research reported in this manuscript is solely the responsibility of the authors, and does not necessarily represent the official views of the National Institutes of Health.

## Author contributions

**Study design:** WLT, AR, RPB, LTG, GHN, CCH and RJMJr. **Experiments and data interpretation:** WLT, AR, RPB, LTG, GHN, NM, JO, CCH and RJMJr. **Manuscript writing, Figures, Tables and Supporting Information:** WLT, AR, RPB, LTG, GHN, NM, JO, CCH and RJMJr.

